# Physics-Informed Neural Network for Mapping Vascular and Tissue Dynamics Using Laser Speckle Contrast Imaging

**DOI:** 10.64898/2026.02.01.702939

**Authors:** Shuying Li, Rockwell Tang, Bingxue Liu, Victoria Krepulec, Will Donovan, David Boas, Xiaojun Cheng, Lei Tian

## Abstract

**Significance:** Quantitatively mapping both cerebral blood flow and tissue dynamics from laser speckle contrast imaging (LSCI) is powerful for studying cerebral blood flow and neurovascular coupling, particularly in the context of stroke. Conventional multi-exposure fitting is slow and difficult to scale. Efficient, physically grounded methods are needed to extract both vascular and tissue dynamic biomarkers from LSCI data.

**Aim:** To develop and validate a physics-informed neural network (PINN) that quantitatively estimates fast (vascular) and slow (tissue-related) speckle decorrelation parameters directly from LSCI measurements without requiring ground-truth labels.

**Approach:** We developed a physics-informed neural network (PINN) to estimate fast (vascular) and slow (tissue-related) speckle decorrelation parameters directly from multi-exposure LSCI data without requiring ground-truth labels. The analytical LSCI model is embedded in the network loss function, enforcing consistency with speckle physics during training. The model operates in a self-supervised manner and performs pixel-wise inference across full-field images. The framework was evaluated using *in vivo* mouse stroke LSCI datasets.

**Results:** The PINN accurately recovered fast decorrelation rates associated with cerebral blood flow and slower dynamics linked to tissue and cellular motion. The parameter maps closely match those from traditional nonlinear fitting, but at orders-of-magnitude higher speed, reducing analysis to under one second per image. It also generalizes to unseen subjects and remains robust under noise.

**Conclusions:** Our approach establishes physics-informed learning as a practical framework for near real-time extraction of vascular and cellular biomarkers from LSCI, enabling longitudinal monitoring of stroke progression and potentially facilitating clinical translation.

## 1 Introduction

Understanding and mapping both blood flow and tissue dynamics in the brain is critically important, especially in the context of stroke where cerebral blood flow (CBF) directly affects tissue dynamics and survival, while changes in cellular activity often accompany vascular events.^1–3^ Monitoring not only perfusion but also intracellular motility can provide early biomarkers of tissue viability, stroke progression, and response to therapy.^4,5^ Laser Speckle Contrast Imaging (LSCI) has emerged as a powerful tool for this task.^4,6–8^ LSCI is a wide-field, high-speed optical imaging technique that quantifies blood flow by analyzing speckle pattern fluctuations: faster motion like red blood cell (RBC) flow leads to reduced speckle contrast. It has been widely used in preclinical studies including murine stroke models to map CBF dynamics across the cortex, and even translated to clinical settings including rheumatology, wound assessment, dermatology, ophthalmology, endoscopy, dentistry, neurology, and intraoperative monitoring.^9–18^ Recent advances showed that LSCI can also capture slow speckle dynamics associated with cellular or tissue motions, distinct from the fast blood flow signals.^4,19^ These slow dynamics provide a window into cellular health including changes in organelle transport, cytoskeletal remodeling, and membrane fluctuations, activities that depend on cellular energy availability.^20–22^ Mapping both the fast and slow components thus offers a more comprehensive picture of tissue status during ischemia stroke, combining vascular blood-flow information and slow tissue dynamics that may reflect cellular-scale motion and metabolic activity.

Traditionally, extracting quantitative blood flow and tissue dynamics information from LSCI has relied on pixel-wise curve fitting of speckle contrast data.^4,8,19,23–31^ In a typical approach, multi-exposure LSCI measurements are made by acquiring speckle images at a fixed frame rate and combining frames to synthesize various camera exposure durations.^26,27^ An analytical model is fitted at each pixel using nonlinear least-squares optimization methods such as Trust-Region^29,32^ or Levenberg–Marquardt algorithm.^26,30^ During fitting, the solver iteratively adjusts model parameters, such as the fast-flow decorrelation time (*τ_c_*_1_) and the slow-dynamics decorrelation time (*τ_c_*_2_) to minimize the residual error between the measured speckle contrast and the model-predicted speckle contrast. While this model-based fitting is effective, it comes with significant limitations in terms of computation speed. Performing nonlinear least-squares fits is slow, and mapping a full image can take hours, making real-time or high-throughput monitoring infeasible. One prior attempt to accelerate parameter estimation for single-exposure SFDI explored a Newton iterative approach combined with GPU implementation; however, this method remained limited for samples with large static components, i.e., scattering contributions from stationary or slowly changing structures.^33^ These fitted parameters are model-derived estimates rather than direct ground-truths, and they may be affected by measurement noise, model assumptions, parameter constraints, calibration of model constants, and residual fitting errors.

An alternative to pixel-wise fitting is data-driven models like neural networks.^32,34,35^ A neural network could learn to output flow and tissue parameters given speckle data, accelerating inference after training. However, supervised neural networks would require a large dataset of LSCI measurements with corresponding ground-truth parameter maps for training. Such ground truth is difficult to obtain experimentally because the available parameter maps are estimates from the same curve-fitting procedure that we aim to replace, and no direct gold-standard measurement is available. Moreover, a black-box network that blindly learns a mapping lacks interpretability in terms of physical meaning.

Physics-Informed Neural Networks (PINNs) offer a compelling solution to these issues. PINNs incorporate physical laws or model equations directly into the training of the neural network,^36^ which means the network can be trained without explicit ground-truth outputs by instead forcing it to satisfy the known physics on the observed data. In our case, the physical model is based on dynamic light scattering and encapsulated by the LSCI speckle contrast model, which is the mathematical relationship between speckle contrast and flow/tissue parameters. By embedding this analytical model into the neural network training, the network learns the inverse mapping (from measured speckle data to parameter values) in a self-supervised manner. This approach combines the strengths of both approaches: the interpretability and scientific grounding of the analytical physics-based model, and the flexibility and speed of neural networks.

In this paper, we present a PINN that learns to map LSCI data to blood flow and tissue dynamics parameters. We demonstrate that our PINN achieves comparable accuracy to traditional pixel-wise fitting but is orders of magnitude faster, enabling the mapping of whole images in seconds instead of hours. We validate the method on *in vivo* LSCI data from mouse stroke experiments, showing that it can track the spatiotemporal evolution of both CBF and slow cellular dynamics during stroke, with the PINN outputs closely matching those of nonlinear curve fitting. The PINN requires no externally labeled training data (only the measured speckle images and the governing equations of speckle contrast), addressing the lack of ground truth data and reducing the risk of non-physical predictions.

## 2 Methods

### 2.1 Laser Speckle Contrast Imaging (LSCI)

Experiments were conducted using a custom LSCI system^4^ (schematized in Fig. 1a). We employed an epi-illumination geometry for enhanced sensitivity to slow dynamics. A collimated 853 nm laser diode (Topica WS) was directed perpendicular to the cortical surface (epi-illumination), illuminating on the mouse cortex through a cranial window. The backscattered light was collected by a 1.5× magnification imaging lens and detected with a CMOS camera (Basler acA1440-220um, 8-bit) placed above the head. A linear polarizer (P1, Thorlabs LPNIR100) before the sample and an orthogonally oriented polarizer (P2, Thorlabs LPNIR100) in front of the camera to primarily pass multiply-scattered light from deeper tissue. The camera was operated at high frame rates to capture raw speckle images. For fast speckle dynamics (FSD) measurements (sensitive to blood flow), we set the exposure time to 4.4 ms, the minimum exposure of our camera corresponding to an ∼224 Hz frame rate.^4^ For slow speckle dynamics (SSD) measurements (sensitive to slower tissue motions), we used a longer exposure of 100 ms, with the camera running at 10 Hz. During imaging, laser power was adjusted by modulating the driving current such that the average raw camera counts were maintained at ∼40 Analog-to-Digital units out of 255 to avoid saturation while keeping shot noise contributions negligible.

**Fig. 1.**
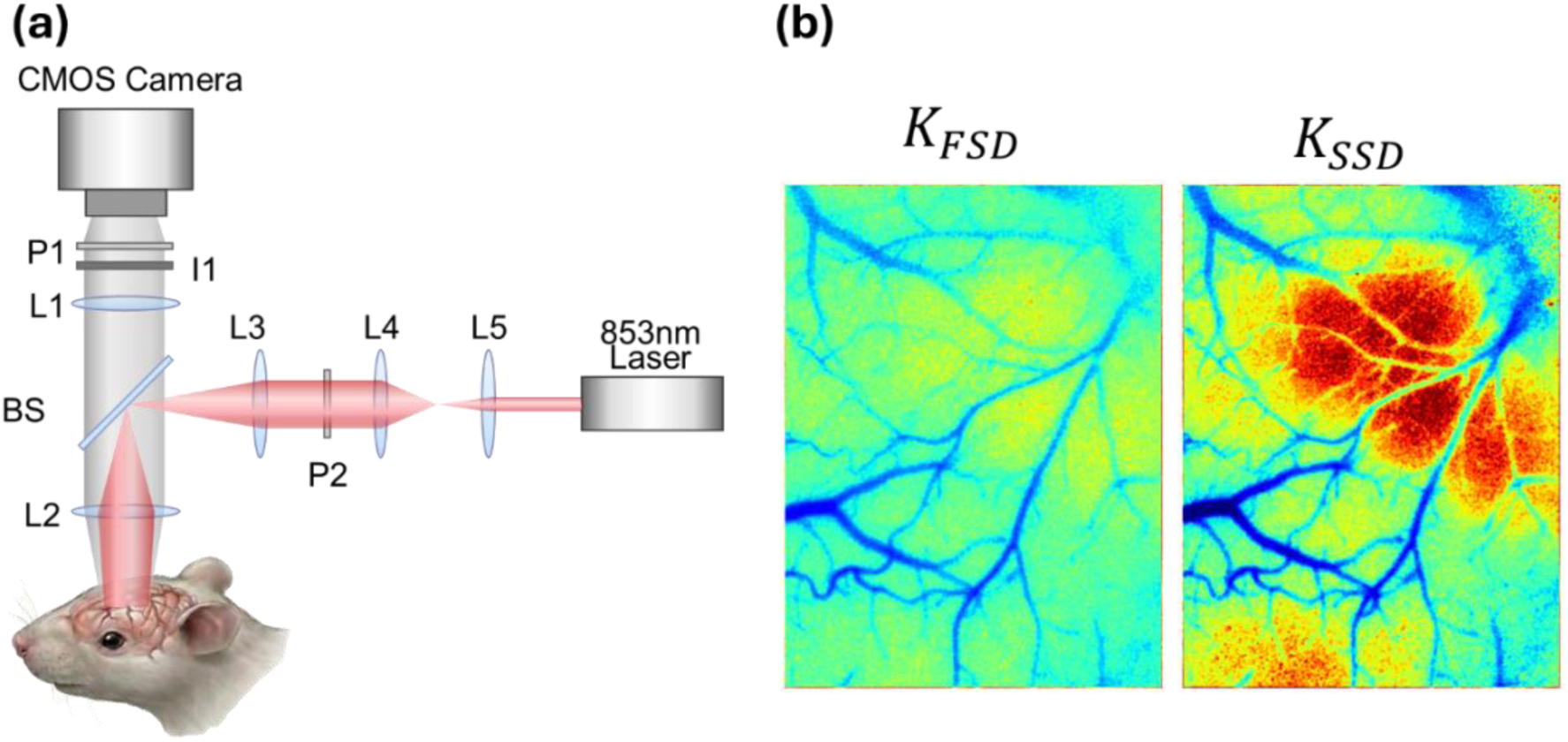
(a) Schematic of the laser speckle contrast imaging (LSCI) setup, recreated from Liu et al.^4^ The mouse head illustration was generated using ChatGPT (version 5.2, OpenAI) for conceptual visualization only. All other graphical elements were created by the authors. P: polarizer. L: lens. BS: beam splitter. I: iris. P1 and P2 are cross-polarized. (b) Examples of speckle contrast map K for fast speckle dynamics (FSD) and slow speckle dynamics (SSD).

Mice were anesthetized with isoflurane (3% for induction and 1–2% for maintenance during imaging) and prepared with a 2-mm-diameter craniotomy in the temporal bone over the distal middle cerebral artery (dMCA) to expose the vessel, as detailed in our prior work.^4^ In the stroke experiments, either a permanent distal middle cerebral artery (dMCA) occlusion was induced by FeCl3-mediated thrombosis, or a transient 90-minute dMCA compression was performed, followed by reperfusion. Imaging sessions included a baseline (pre-stroke), acute stroke, and several post-stroke intervals, with separate acquisitions for FSD and SSD at each time point. A spectral-domain optical coherence tomography (OCT) system with a 1310 nm center wavelength and 170 nm bandwidth (Thorlabs) was used to identify the stroke core, following previously reported acquisition parameters and processing methods.^4,37,38^ All experiments and animal procedures were approved by the Boston University Institutional Animal Care and Use Committee.

### 2.2 Theory and model fitting for fast and slow dynamics

The raw speckle images were processed to compute the spatial speckle contrast, defined as:^4^

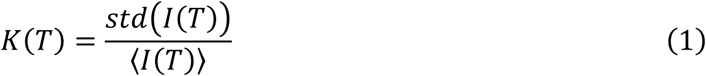

where *I*(*T*) is the raw intensity image acquired with exposure time *T*, *std* denotes the standard deviation of pixel intensities in a local window (here 7×7 pixels), and ⟨*I*(*T*)⟩ is the mean intensity in that window. To capture the dependence of *K*(*T*) on exposure time, we utilized a multi-exposure strategy via frame binning. Consecutive high-frame-rate images were numerically summed (integrated) to emulate longer exposures post hoc.^4,26,27^ This synthetic-exposure approach is justified because camera exposure represents temporal integration of detected intensity over the exposure duration. Therefore, summing adjacent short-exposure frames approximates the intensity that would have been accumulated during a single longer exposure, provided that illumination is stable, the camera response is linear, and interframe delays are sufficiently short relative to the speckle decorrelation time. ^27^ At longer exposure times, total detected intensity increases relative to shorter exposures if illumination power is unchanged, which can alter the shot-noise contribution to measured speckle contrast. However, because shot-noise-related bias is inversely proportional to photon flux per speckle, camera quantum efficiency, and exposure time,^39^ this bias is expected to be small under the high-photon-flux conditions of our mouse brain recordings using a modern high-quantum-efficiency camera. We acquired 10 s of FSD data and 5–10 min of SSD data and averaged the contrast over time to get a low-noise estimate of *K*(*T*) at each exposure duration. This approach yields a contrast vs. exposure time curve for every pixel.

We employ a two-component speckle model reflecting fast blood flow and slow tissue motions.^4^ The fundamental assumption is that dynamic scattering in tissue comes from two populations of moving scatterers^4,19^: (1) RBCs (fast) with a short decorrelation time *τ_c_*_1_ (on the order of a few milliseconds), and (2) intracellular and subcellular motions, such as organelle transport, cytoskeletal remodeling, and membrane fluctuations (slow) with a much longer decorrelation time *τ_c_*_2_ (order of seconds)^22^. The detailed development of this model has been described in prior work.^4,19,31,40–46^ For short exposure times (*T* ≪ *τ_c_*_2_), the slow component is essentially static during the exposure, so the contrast is dominated by fast dynamics. In this regime we use a model *K_FSD_*(*T*) that includes *τ_c_*_1_ and *ρ*_1_ as parameters (while slow dynamics are negligible):

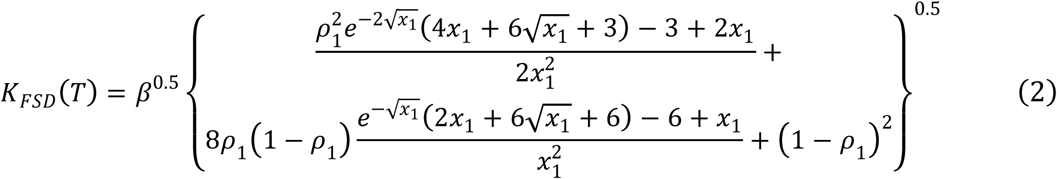

where *β* accounts for loss of coherence due to speckle averaging, polarization, instability, or source incoherence; *x*_1_ = *T*/*τ_c_*_1_; *τ_c_*_1_ is the FSD decorrelation time constant; *ρ*_1_ is the average fraction of the dynamic component for FSD. We define *FSD rate* = 1/*τ_c_*_1_ (larger *τ_c_*_1_ means slower blood–flow dynamics).

Conversely, for long exposure times (*T* ≫ *τ_c_*_1_), the fast RBC-induced speckle fluctuations have fully decorrelated (RBC speckles blur out within the exposure), and the remaining contrast is governed by the slow dynamics with parameters *τ_c_*_2_ and *ρ*_2_. We denote this long-exposure model *K_SSD_*(*T*):

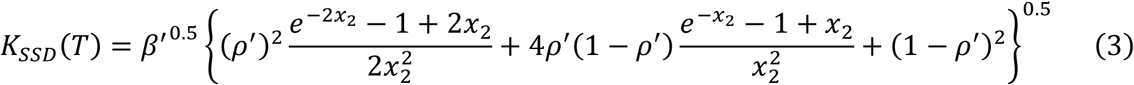

where *x*_2_ = *T*/*τ_c_*_2_; *ρ*_2_ is the average fraction of the dynamic component for SSD, i.e., the fraction of detected photons scattered by slowly moving or deforming tissue cells; *β*′ = *β*(1 − *ρ*_1_)², and *ρ*′ = *ρ*_2_/(1 − *ρ*_1_). We define *SSD rate* = 1/*τ_c_*_2_ (larger *τ_c_*_2_ indicates slower cellular/tissue dynamics).

In traditional curve fitting, we fit the measured contrast curves in two segments: FSD data (*T* from 4.4 ms up to ∼440 ms) were fit with the fast model to extract *τ_c_*_1_ and *ρ*_1_, and SSD data (*T* from 1 s up to 40 s) were fit with the slow model to extract *τ_c_*_2_ and *ρ*_2_. This split fitting mitigates interference between the two processes and is possible because of the two to three order magnitude difference in *τ_c_*_1_ and *τ_c_*_2_. The output is four parametric maps: two blood flow maps (fast decorrelation time *τ_c_*_1_, and the dynamic fraction *ρ*_1_) and two tissue dynamics maps (slow decorrelation time *τ_c_*_2_, and fraction *ρ*’).

Data were fit to the models with custom scripts using MATLAB’s `fit` function. For FSD (Eq. 2), *β* was fixed at 0.72, as determined from the static phantom measurement, to reduce the number of free fitting parameters.^47^ For SSD (Eq. 3), *β*′ was set to the mean spatial contrast *K*(*T*_0_)^2^ at the first fitting timepoint. The dynamic fractions *ρ*_1_ and *ρ*′ were obtained from fitting constrained to the range [0,1]. Fits were performed as nonlinear least-squares optimizations using MATLAB’s default Trust-Region algorithm. Robust least absolute residual fitting (`Robust = ’LAR’`) was used to reduce sensitivity to outliers. The pixel-wise nonlinear curve-fitting calculations were performed with 28 CPU cores with 8GB memory per core on Boston University’s shared high-performance computing cluster (BU SCC), which includes a heterogeneous CPU infrastructure with multiple CPU types depending on compute nodes.

For computational speed comparison purposes, GPU-accelerated robust nonlinear fitting was implemented in MATLAB using a customized Trust-Region-Reflective algorithm applied simultaneously to all image pixels. LAR robust fitting was achieved via iteratively reweighted least squares, with residual-based weights updated at each iteration. At each step, a Gauss–Newton update was computed from the weighted normal equations and constrained by a trust-region bound. Step acceptance was determined using the ratio of actual to predicted reduction in the objective function, and the trust-region radius was adjusted adaptively based on this ratio. Bound constraints on parameters were enforced via projection onto the feasible interval after each update. All forward-model evaluations, Jacobian computations, linear solves, and parameter updates across pixels were performed simultaneously using MATLAB `gpuArray` and `pagefun`. The GPU and CPU implementations shared the same physical model, parameter bounds, and convergence tolerance. The two implementations may not be identical as the CPU method used MATLAB Curve Fitting Toolbox’s proprietary robust fitting procedure.

### 2.3 Physics-Informed Neural Network (PINN)

The physics-informed neural network is illustrated in Fig. 2(c). The role of this network is to take the place of pixel-wise curve fitting by learning the mapping from a pixel’s speckle contrast measurements to that pixel’s underlying parameters. Compared to conventional least-squares fitting shown in Fig. 2(a), PINN optimizes the neural network weights globally based on the measurements from all pixels, instead of fitting each pixel independently. In addition, rather than training the network with ground truth parameter maps (which we lack) as shown in Fig. 2(b), we train it by requiring consistency with the physical speckle model and the observed measurements. The training data consist of the measured speckle contrast values *K*(*T*) for each pixel at various exposure times. Separate neural networks with the same architecture were trained for the FSD and SSD, using *K_FSD_*(*T*) and *K_SSD_*(*T*) as their input, respectively. These networks serve as inverse models that map measured speckle contrast curves *K_meas_*(*T*) to the corresponding model parameters. The output of the FSD neural network is [*ρ*_1_, *τ_c_*_1_], while the output of the SSD neural network is [ *ρ*′, *τ_c_*_2_]. These predicted parameters are then passed through the corresponding analytical forward FSD and SSD model (Eq. 2 and 3 in Section 2.1) to reconstruct the model-predicted speckle contrast *K_pred_* (*T*).

**Fig. 2.**
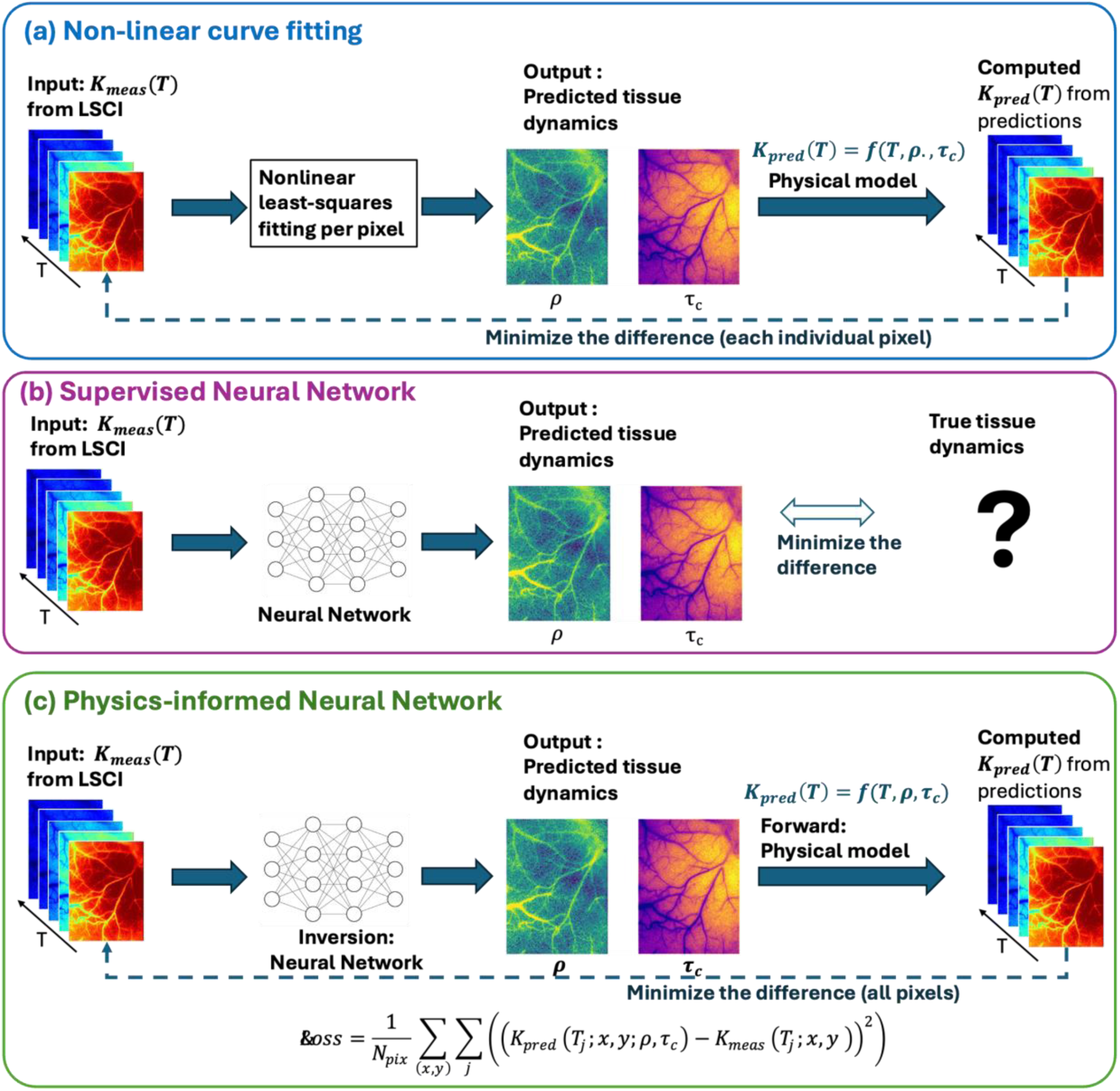
LSCI tissue dynamics reconstruction with (a) Traditional curve fitting (b) Traditional supervised neural network and (c) PINN. Separate neural networks with the same architecture were trained for the FSD and SSD, using *K_FSD_*(*T*) and *K_SSD_*(*T*) as their input, respectively. These networks serve as inverse models that map measured speckle contrast curves *K_meas_* to the corresponding model parameters, [*ρ*_1_, *τ_c_*_1_] for FSD and [*ρ*′, *τ_c_*_2_], for SSD. These predicted parameters are then passed through the corresponding analytical forward FSD and SSD model (Eq. 2 and 3 in Section 2.1) to reconstruct the model-predicted speckle contrast *K_pred_*(*T*). The loss is the MSE difference between *K_pred_*(*T*) and *K_meas_* (*T*), calculated over all pixels and exposure times.

We formulate a loss that penalizes the difference between *K_pred_*(*T*) (using the network’s output dynamics parameters) and the actual measured contrast *K_meas_*(*T*). Specifically, for each pixel and each exposure time *T_j_* in our dataset, we compute a predicted contrast, which is essentially the same form as the FSD and SSD model. We then define the data fidelity loss as the mean squared error (MSE) between the calculated *K_pred_* and the measured *K_meas_* over pixels and exposure times:

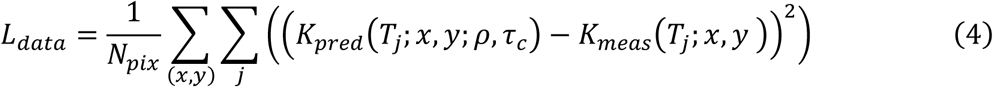

where *T_j_* is the exposure time, *x*, *y* are spatial coordinates, *ρ*, *τ_c_* are the parameters predicted from the neural network. This loss drives the network to produce parameters that reconstruct the speckle contrast curves for every pixel without the need for ground truth *ρ*_1_ and *τ_c_*_1_ or *ρ*′ and *τ_c_*_2_. Because the network is shared across all pixels, it is fitting all pixel curves simultaneously, which allows it to generalize and denoise the solutions by leveraging common patterns in the data that cannot be achieved with traditional pixel-wise fitting.

In addition to the data fidelity term (Eq. 4), we include regularization constraints in the final layer of the neural network to enforce known physical bounds on the parameters. We add Sigmoid to *ρ*_1_ and *ρ*′to limit their range to [0,1], and Softplus to *τ_c_*_1_and *τ_c_*_2_ so they are guaranteed to be non-negative.

#### Data Splits

We compiled a dataset from three mice to train and evaluate the PINN. From these sessions we extracted the contrast-vs-time curves for every pixel under both FSD and SSD acquisitions. This provided a diverse set of speckle curves reflecting different physiological states (normal, ischemia, reperfusion, recovery). Data from Mouse #1 were used for model training and validation. Eight time points were included in the training set: baseline, two acute stroke time points, 3 h, 4 h, 5 h, 1 d, and 2 d post-stroke. Each time point consisted of images with a spatial resolution of 1440 × 1080 pixels, yielding a total of 12,441,600 pixel-wise training samples under the assumption that each pixel is independent. We reserved two intermediate time points from Mouse #1 (specifically, an acute stroke frame and 7 h after stroke in the transient stroke series) as a validation set to tune hyperparameters and we monitored the loss on this held-out condition to avoid overfitting. Mice #2 and #3 (used for testing) underwent a similar stroke imaging protocol but was not used in training at all. These testing data are solely used for evaluating how well the trained PINN generalizes to an unseen subject. All results presented in Section 3 are on these independent test mice to demonstrate generalization.

#### Implementation

We implemented the PINN in Python using PyTorch. The model consisted of three fully connected layers with 128, 64, and 2 neurons, respectively, and rectified linear unit activation function (ReLU) activations applied to the first two layers. The network input dimension was 28, corresponding to the extracted measurement features. To enforce physical plausibility, we applied two output activation functions: the sigmoid function, defined as 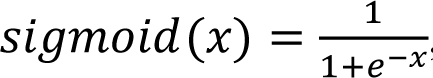, which maps real-valued inputs to the range (0, 1), and the softplus function, defined as *softplus*(*x*) = *ln*(1 + *e^x^*), which maps real-valued inputs to positive values. Specifically, *ρ*_1_or *ρ*’ was passed through a sigmoid activation to limit its range to (0, 1), and *τ_c_*_1_ or *τ_c_*_2_ was passed through a softplus function to ensure positive values. The two constrained outputs were concatenated to form the final prediction vector [*ρ*_1_, *τ_c_*_1_] or [*ρ*′, *τ_c_*_2_]. The network was trained using the Rprop optimizer (learning rate 1e−2) for 200 epochs, with batch size = 32768. Rprop updates each parameter based on the sign of the gradient rather than its magnitude, with adaptive parameter-specific step sizes that improve robustness to gradient scaling. Importantly, the PINN training consumed the raw measured data only and the known equations, and no ground-truth parameter maps were needed anywhere in this pipeline. The final trained model was then used to predict *ρ* and *τ_c_* for each pixel in the test mouse images (inference step). This simply involves a forward pass of the network for each pixel coordinate, which is highly efficient. We also computed the conventional fitted parameters for the test mouse (using the dual-exposure fitting described in Liu et al. ^4^) to serve as a reference for accuracy and to mimic “ground truth” for comparison, acknowledging that these fitted values themselves have some uncertainty.

#### Evaluation metrics

To quantitatively assess the performance of the proposed models, we compared the speckle contrast *K_pred_*(*T*) computed from PINN predicted parameters, with the experimental measurements *K_meas_*(*T*). Performance was evaluated using three metrics: Mean Absolute Error (MAE), Mean squared error (MSE), and the coefficient of determination (*R*²), which is computed as:

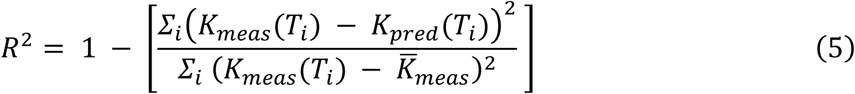

where *K_meas_*(*T_i_*) denotes the measured speckle contrast at exposure time *T_i_*, *K_pred_* (*T_i_*)is the speckle contrast reconstructed from the model-predicted parameters at the same exposure time, and *K̅_meas_* is the mean of the measured speckle contrast across all exposure times. *R*² quantifies the proportion of variance in the measured speckle contrast explained by the model, indicating overall fitting quality.

Statistical analysis between PINN and curve fitting results was performed using the Wilcoxon signed-rank test. Differences with *p* < 0.05 were considered statistically significant.

## 3 Results

### 3.1 Qualitative Results

#### Examples of predicted tissue dynamics maps

The physics-informed neural network successfully predicted high-quality maps of the tissue dynamics parameters on the unseen test mice, closely matching the results of traditional curve fitting. Figures. 3 and 4 compare the PINN-predicted maps of fast and slow speckle parameters to those obtained by CPU-based nonlinear least-squares fitting for representative cases. In these figures, we show baseline (pre-stroke), stroke, and post-stroke (2 days after a permanent MCA occlusion) time points for Mouse #2. Each panel displays the spatial map of the fast and slow dynamics decorrelation time constant (*τ_c_*_1_ and *τ_c_*_2_) and the average fraction of the dynamic component for FSD and SSD (*ρ*_1_ and *ρ*′) as computed by the PINN vs. by pixel-wise curve fitting. Visually, the PINN and conventional maps are almost indistinguishable. Both methods reveal the same anatomical and physiological patterns. We see that within the stroke core, *τ_c_*_1_ and *τ_c_*_2_ both increase, indicating slower blood flow and cellular dynamics, while *ρ*_1_ the fraction of FSD decreases. Figures 3 and 4 also show spatial maps of the coefficient of determination *R*^2^, with nearly all pixels exhibiting values close to 1.0 (green). Slightly lower *R*^2^ values are observed along blood vessel regions but remain above 0.95, indicating strong local consistency between the measurements reconstructed from the predicted parameters and the true measurements.

**Fig. 3.**
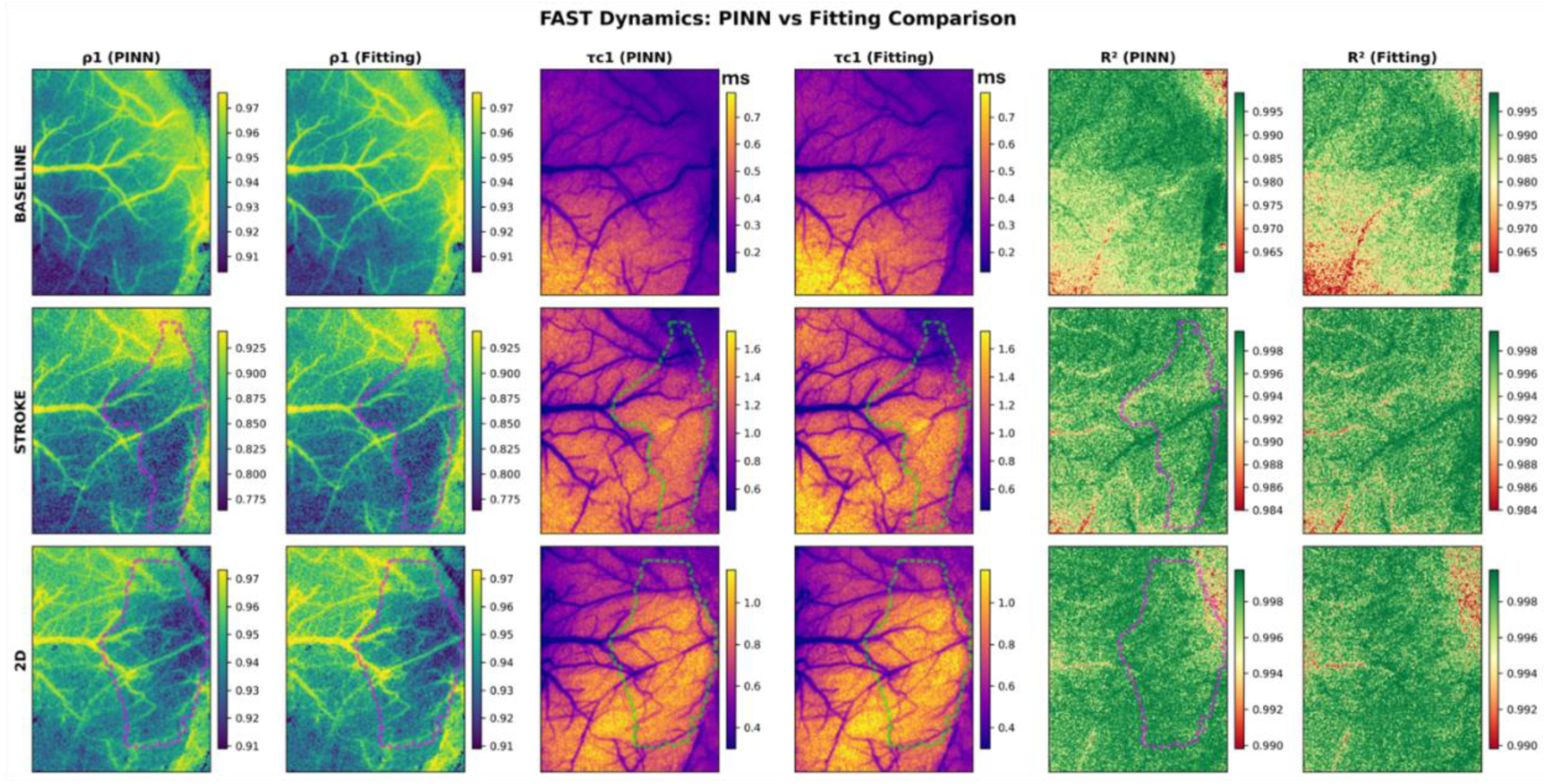
Comparison of PINN-predicted and conventionally fitted fast dynamics parameters on the test mouse. Dashed lines indicate the stroke core, as determined by optical coherence tomography (OCT).^4^

**Fig. 4.**
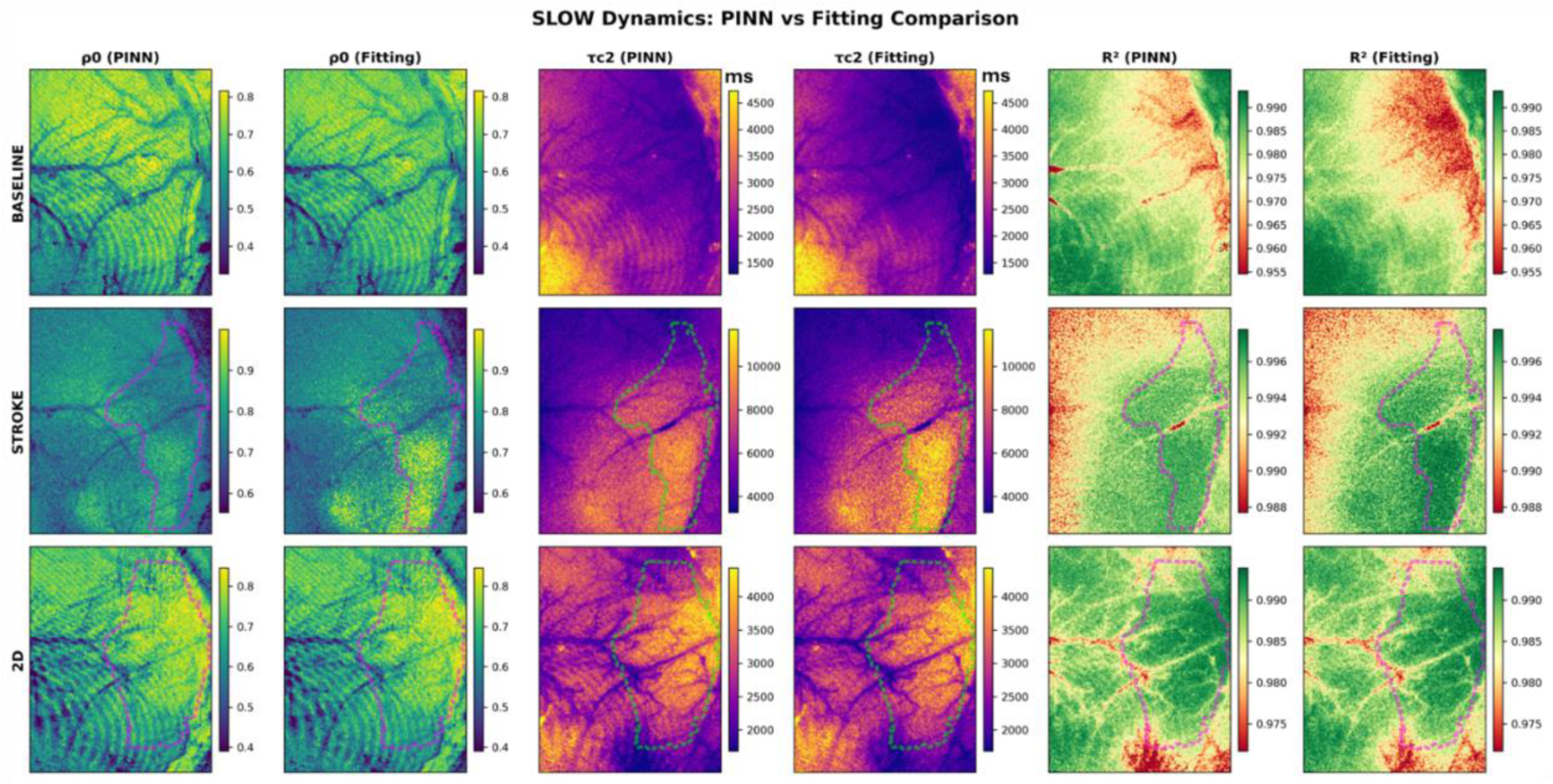
Comparison of PINN-predicted and conventionally fitted slow dynamics parameters on the test mouse, the same as that in Fig. 3.

#### Time-Series Tracking of Stroke

We further examined how well the PINN could track the evolution of blood flow and tissue dynamics over time in the test mouse, compared to conventional analysis. Figure 5 presents a time series of parametric maps for Mouse #2 (permanent stroke model) at eight key time points: baseline, acute stroke, 3 h, 4 h, 5 h, 7 h, 1 d, and 2 d post-stroke. We show maps of the fast decorrelation rate (1/*τ_c_*_1_, proxy for CBF) and slow decorrelation rate (1/*τ_c_*_2_) as generated by the PINN vs. by curve fitting. Once again, the PINN results mirror the conventional results at each time. For blood flow, both methods show the immediate drop in 1/*τ_c_*_1_ in the occluded region, and this low flow persists through 2 d. For the slow dynamics, both methods show a distinct temporal pattern: during stroke, 1/*τ_c_*_2_ is sharply reduced in a broad cortical area around the stroke core, indicating a substantial slowing intracellular dynamic in the early acute phase. After 3 h post-stroke, the region of slowed dynamics contracts and 1/*τ_c_*_2_ begins to rise again. At 24–48 h, the 1/*τ_c_*_2_in the core remains lower than baseline but higher than the stroke. This initial depression of speckle dynamics followed by partial recovery was highlighted in the original study as a potential indicator of evolving cellular activities.^4^ Our PINN results identically reproduce these spatiotemporal trends, as seen by the close correspondence with the fitted maps at each time point in Fig. 5. This confirms that the PINN can be trusted to monitor dynamics longitudinally. These qualitative comparisons demonstrate that the PINN is not only producing plausible-looking maps, but it is essentially recovering the same information as the time-tested non-linear fitting technique.

**Fig. 5.**
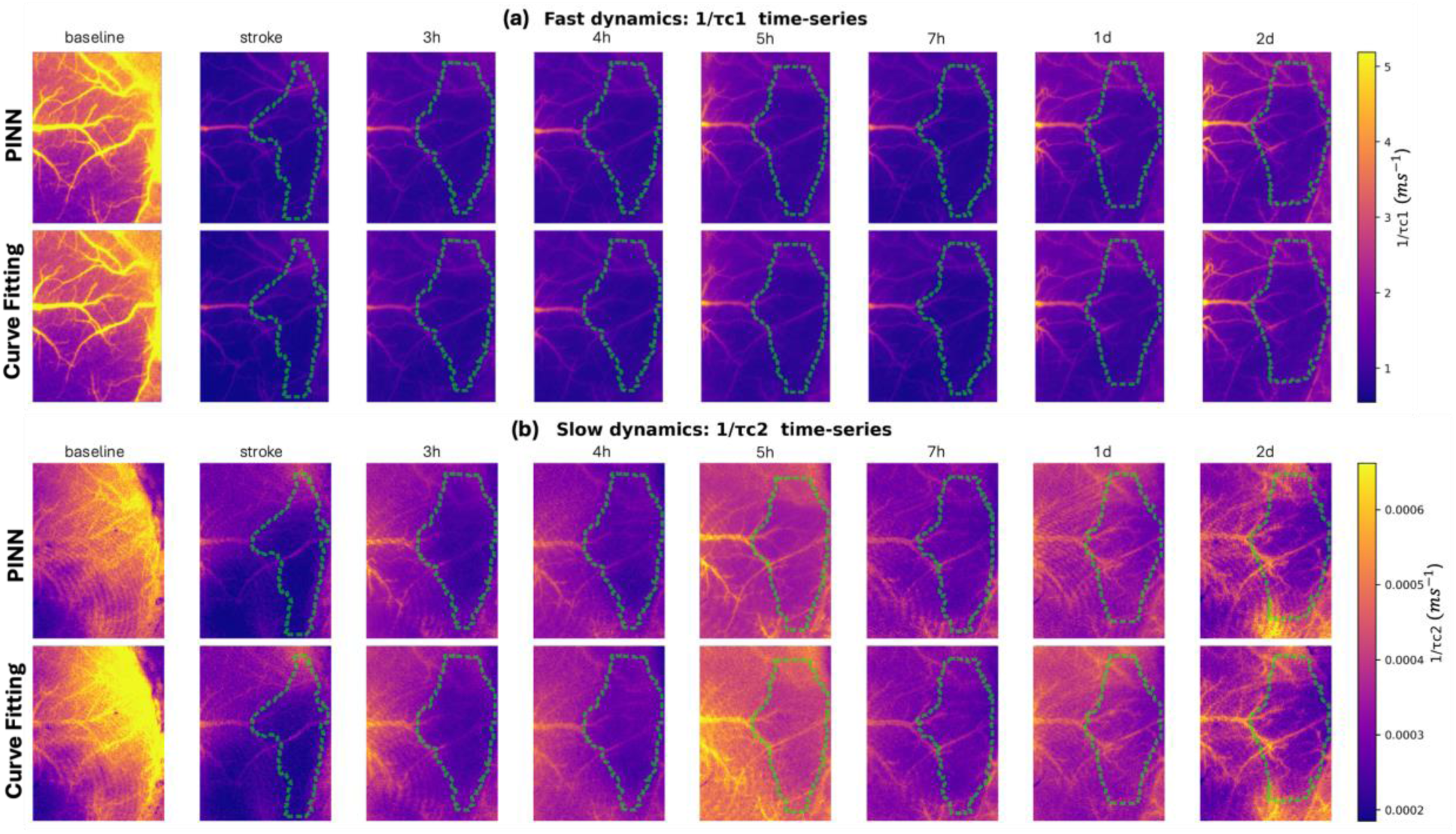
Time course of stroke progression in the permanent occlusion model, as seen through fast and slow speckle dynamics (test mouse #2). Maps of (a) blood flow (FSD rate 1/*τ_c_*_1_) and (b) slow tissue dynamics (SSD rate 1/*τ_c_*_2_) are shown at baseline and various times post-stroke, for both conventional fitting and PINN. Dashed lines indicate the stroke core.

### 3.2 Quantitative Evaluation

We quantitatively evaluated the PINN’s performance against the traditional curve-fitting approach in terms of accuracy, computational efficiency, and noise robustness.

#### Accuracy

Quantitative evaluation was performed by comparing the reconstructed speckle contrast curves *K_pred_* (*T*) obtained from the PINN-derived parameters and from nonlinear curve fitting against the true measured *K_meas_*(*T*) for all testing images (*n* = 44), as shown in Fig. 6. Each data point is the averaged value of all pixels for one 1440 × 1080 map. The PINN achieved a similar mean absolute error (MAE, *p* = 0.940) but demonstrated significantly higher coefficient of determination (R², *p* < 0.05) and lower mean squared error (MSE, *p* < 0.001) compared to the fitting baseline. Compared with CPU fitting, the PINN achieved a similar mean absolute error (MAE, *p* = 0.940) but demonstrated significantly higher coefficient of determination (R², *p* < 0.05) and lower mean squared error (MSE, *p* < 0.001). Compared with GPU fitting, the PINN showed similar R² and MSE (*p* > 0.05), but a higher MAE (PINN: 0.00180; GPU: 0.00151; *p* < 0.001). This slightly higher MAE may be partly attributable to the MSE-type loss function used during PINN training. The nonlinear fitting baseline used robust LAR weighting, which is more closely aligned with the MAE criterion. Overall, the PINN reconstructed the measured speckle contrast curves with accuracy comparable to that of conventional fitting methods.

**Fig. 6.**
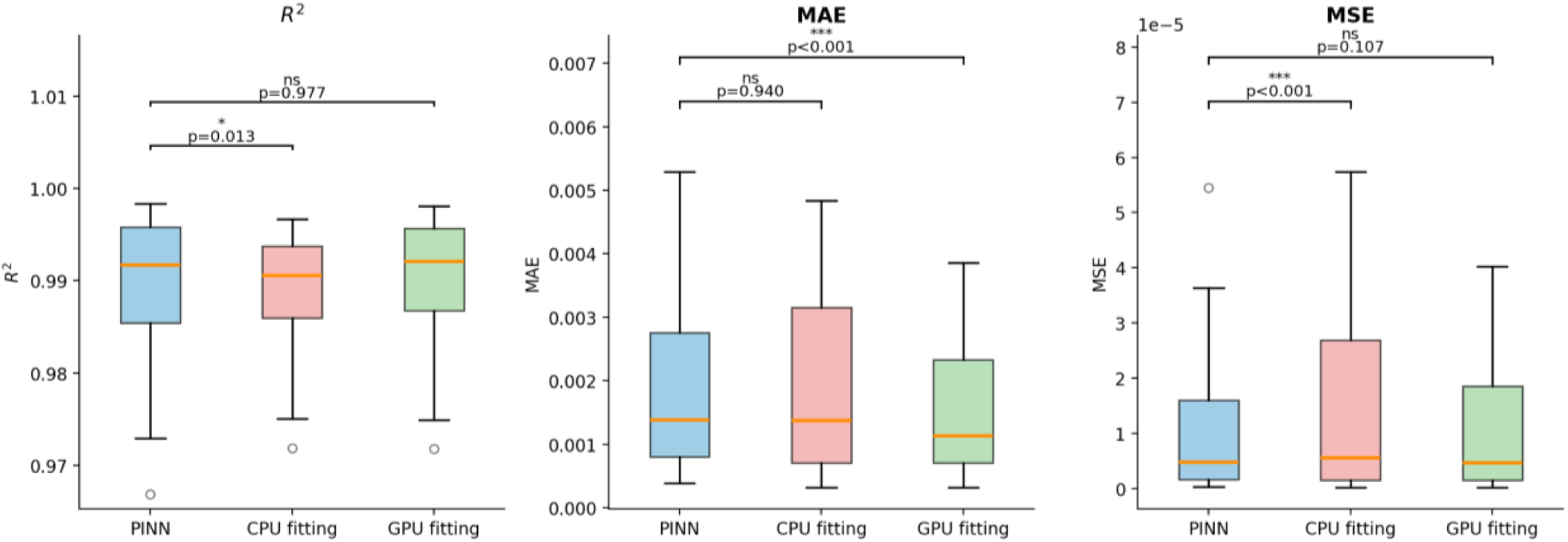
Quantitative performance comparison of PINN vs. traditional fitting for all testing images (*n* = 44). The comparison is between the reconstructed speckle contrast *K_pred_*(*T*) obtained from the PINN-derived parameters and from nonlinear curve fitting against the true measured *K_meas_*(*T*).

#### Computational Speed

The PINN approach offers a tremendous speed advantage. The training for each model takes about 11 hours on a NVIDIA A40 GPU. After training the model (which is done offline), applying it to new data (inference) is extremely fast. For a 1440 × 1080-pixel image, the trained PINN generated both fast and slow tissue dynamics maps in about 0.4s of computation time (excluding file I/O and including CPU-GPU data transfer) on an NVIDIA RTX3090 GPU, demonstrating its efficiency for near–real-time analysis. This processing speed is compatible with near-real-time analysis and could support real-time implementation if integrated with an optimized acquisition and preprocessing pipeline. In contrast, traditional pixel-wise fitting required approximately 1–1.5 hours to process the same image on 28 CPU cores using parallelized MATLAB implementations, and approximately 1.5–3 minutes on an NVIDIA RTX3090 with GPU implementations. Overall, the PINN approach achieved approximately two to four orders of magnitude faster speed than traditional fitting methods. This speed up becomes even more important in time-series imaging: for example, processing 10 time points with curve fitting might take ∼10–15 hours on a CPU and 15-30 minutes on a GPU, whereas the PINN can process all 10 within seconds. In practice, this means one could envision the PINN being deployed during an imaging session to continuously display updated blood flow and tissue status maps on the fly, which was not realistic with previous computational methods. The light weight of the PINN also suggests it could run on embedded systems or GPUs in the imaging device for point-of-care applications.

#### Robustness to Noise

We simulated fast and slow speckle dynamics using analytical forward models parameterized by *τ_c_*_1_, *τ_c_*_2_, *ρ*_1_, and *ρ*′ values sampled from physiologically realistic ranges: *τ_c_*_1_ was varied from 0.2 to 1.8 ms, *τ_c_*_2_ from 2,000 ms to 12,000 ms, *ρ*₁ from 0.7 to 1, and *ρ*′ from 0.3 to 1, with *β* fixed at 0.72 and *β*′ fixed at 0.01. Each parameter was discretized into 20 uniformly spaced values, and parameter combinations were used to generate speckle contrast curves *K*(*T*) using Equations (2) and (3). The trained PINN was then evaluated on these synthetic datasets perturbed by additive Gaussian noise ranging from 0% to 10%. For each noise level, the models predicted parameter values from noisy *K*(*T*) inputs. We quantified robustness by computing mean squared error (MSE) and mean absolute error (MAE) between predicted and ground-truth parameters across all samples. The results were aggregated across noise levels for both dynamic regimes, as shown in Fig. 7. Both MAE and MSE remain stable up to 10% noise, confirming robustness of the PINN model in simulation.

**Fig. 7.**
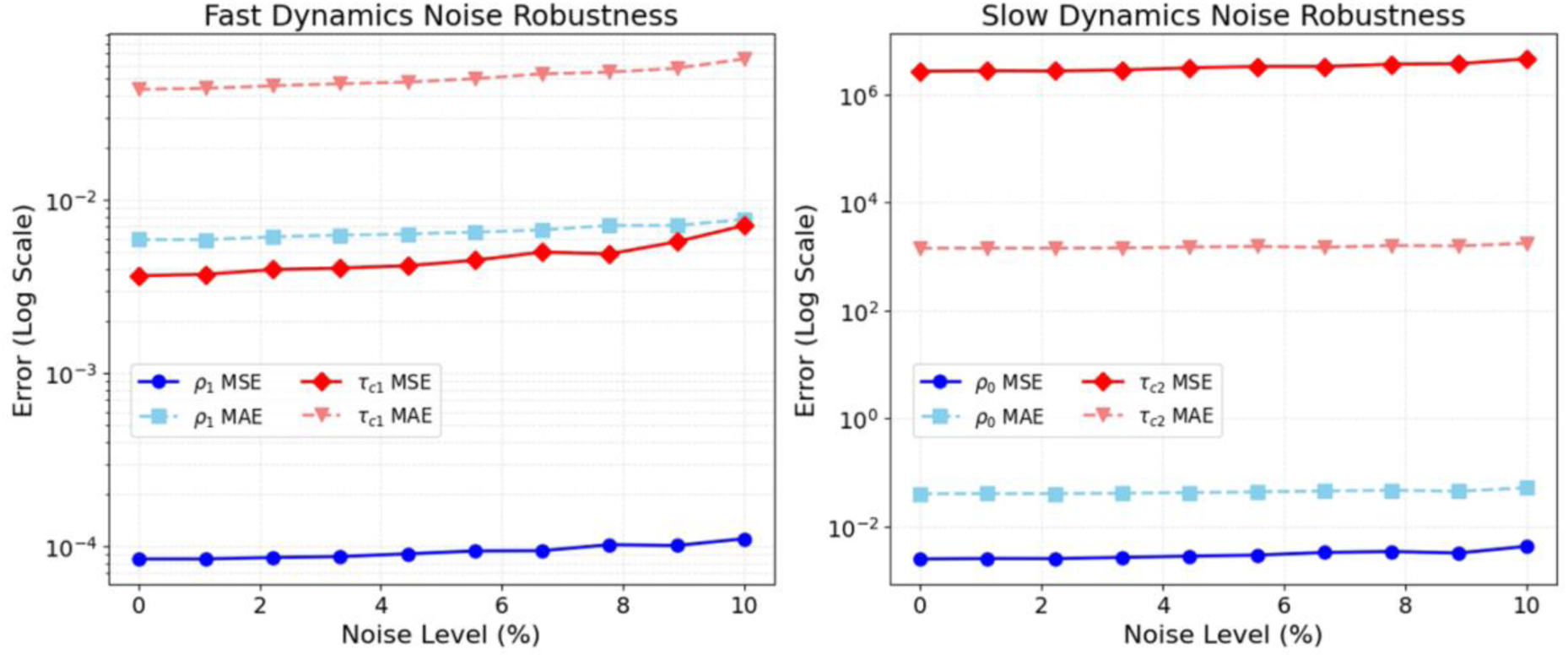
PINN estimation errors as a function of increasing Gaussian noise for simulated data.

## 4 Discussion and Conclusion

We have demonstrated a physics-informed neural network approach for mapping blood flow and tissue dynamics from LSCI data and shown that it can achieve comparable accuracy to the conventional pixel-wise analytical fitting, but with orders-of-magnitude faster computation. To our knowledge, this is the first application of PINNs in the context of LSCI reconstruction. Because the PINN is constrained by the same analytical speckle contrast models used in conventional curve fitting, it estimates the same model-derived parameters rather than discovering new physiological biomarkers. Therefore, the primary contribution of this work is computational acceleration and improved practical usability of quantitative real-time LSCI analysis, while preserving the physical interpretability of the conventional model-based framework.

The PINN was able to accurately recover the fast and slow speckle correlation times (*τ_c_*_1_and *τ_c_*_2_) and the relative contributions of each dynamic component (*ρ*_1_, and *ρ*′), thus providing both vascular (CBF-related) and cellular dynamical information from the LSCI data. This was achieved without needing any ground-truth maps for training because the network learned directly from the measured speckle contrasts by enforcing the well-established speckle contrast theory in its loss function. This approach addresses several limitations in the LSCI quantification workflow. First, it eliminates the heavy computational burden of iterative curve fitting, paving the way for real-time imaging of both blood flow and tissue dynamics. This is especially relevant for stroke or other acute conditions where timely information is critical; with the PINN, one could feasibly monitor the tissue viability changes or assess therapy effects on the fly. The ability to get real-time feedback on both perfusion and cellular activity could guide interventions. For instance, in stroke therapy, a rapid increase in *τ_c_*_2_ could indicate impending tissue infarction even if blood flow partially recovers, consistent with evidence that infarct growth can occur despite successful endovascular reperfusion.^48,49^ Second, the PINN’s physics-based nature ensures that its outputs remain interpretable and physiologically grounded. Unlike a black-box neural network, our PINN inherently respects the relationships defined by the speckle model. This interpretability is a major advantage when introducing AI methods into biomedical research since it helps build trust that the model is not hallucinating features but rather following known physics.

The network was trained exclusively on data from Mouse #1. All the results reported in the paper were testing results obtained from Mice #2 and #3, which were never used during training or validation. The trained PINN generalized to these unseen test mice across all imaging time points, yielding accurate parameter maps without any fine-tuning. Although the PINN was trained using data from a single mouse, each pixel was treated as an independent training sample. Across the wide field of view, pixels different types of tissues and therefore exhibit a broad range of speckle decorrelation behaviors associated with different flow speeds and tissue dynamics. As a result, even data from a single animal provide a large and diverse set of physically distinct speckle contrast curves. In essence, the PINN learned a physics-constrained inverse mapping from measurements to tissue parameters and the range of plausible parameter values, so it could infer parameters for new data that follow the same physical principles and fall within a similar range of optical and physiological conditions represented in the training data. This suggests that the PINN model could be re-used across comparable experiments to obtain the same parametric insights without spending hours in data analysis or worrying about initial guesses for fits.

Building on this work, several extensions can be explored. 1) Enhanced robustness: While our PINN already showed some robustness to noise, further improvements are possible. For example, implementing a realistic noise-aware PINN training where the loss function allows some tolerance to noise could make the model even more resilient. Additionally, including more a priori knowledge like spatial smoothness of parameters as additional physics constraints could regularize the solutions in low-SNR scenarios. 2) Joint fast/slow dynamics modeling: In this study, we treated fast (CBF) and slow (cellular) dynamics separately, in line with the experimental design of dual exposures. An ambitious next step is to create a unified PINN that simultaneously recover both *τ_c_*_1_ and *τ_c_*_2_ in one network. 3) Broader applicability and clinical translation: So far, we demonstrated the PINN on mouse cerebral cortex data. It would be valuable to test the approach on other LSCI applications like monitoring blood flow and tissue viability in skin flaps, or in intraoperative settings where LSCI is used. The framework may be reusable when the underlying speckle contrast model, imaging configuration, and parameter ranges remain comparable. However, because the network was trained using data from a single mouse brain, it may not fully capture variability in blood flow, tissue viability, tissue scattering properties, or physiological conditions across other animal models, humans, or organs beyond the brain. Therefore, we do not assume comparable accuracy without additional validation. If the pretrained mouse-brain model does not generalize well to a new application, the same framework could be retrained or fine-tuned using measured speckle contrast data from the target setting. When applicable, this approach could provide the same model-derived parametric information while reducing the need for time-consuming pixel-wise nonlinear fitting and sensitivity to initial parameter guesses. In that case, the PINN can be retrained or fine-tuned on those contexts by plugging in the appropriate physical models which might include different scattering geometries or additional dynamics. 4) Validation using photon-counting method: Experimentally measured ground-truth parameter maps were not available in this study. However, related decorrelation or blood-flow parameters can be estimated using independent photon-counting approaches, including SPAD-based methods^50,51^ that measure temporal speckle fluctuations and fit autocorrelation functions to estimate flow-related dynamics. These approaches may therefore provide important independent validation data in future studies.

In conclusion, we presented a PINN framework that learns the mapping from LSCI data to underlying tissue dynamic parameters by incorporating the known speckle physics into the network’s training. The approach resolves the limitations of traditional pixel-wise curve fitting by drastically reducing computation time from hours to seconds without the need for ground truth labels. The PINN’s outputs in mouse stroke models closely matched those from conventional curve-fitting, successfully tracking both cerebral blood flow and slow cellular dynamics over time. This highlights the promise of PINN in biomedical imaging. It allows us to retain physical interpretability and accuracy while harnessing the efficiency of neural networks. Going forward, such models could be widely applied wherever we have a forward model but face an intractable inverse problem. Our work enables real-time, label-free mapping of both vascular and cellular health indicators in tissue, which could open new avenues for monitoring strokes and other pathologies with unprecedented speed and detail.

## Disclosure

The authors declare that there are no financial interests, commercial affiliations, or other potential conflicts of interest that could have influenced the objectivity of this research or the writing of this paper.

## Acknowledgments

The authors acknowledge funding from National Institutes of Health (R01EB034272, R01NS127156), a grant from 5022 - Chan Zuckerberg Initiative DAF, an advised fund of Silicon Valley Community Foundation. We also thank Po-kai Su for assistance in GitHub code release. During the preparation of this manuscript, ChatGPT (OpenAI) was used to expand and edit the text written by the authors.

## Code and Data Availability

The data used for model training and testing were generated as part of previous studies and are publicly available through the Distributed Archives for Neurophysiology Data Integration (DANDI) at https://dandiarchive.org/dandiset/001541. The trained model checkpoints and associated training and testing code, along with GPU and CPU implementation of the nonlinear curve fitting, have been made publicly available at are publicly available on GitHub at https://github.com/MiamiU-Li-lab/LSCI_PINN_original.

